# FcγRI is the key determinant of antibody-mediated Zika virus infection of human placental macrophages

**DOI:** 10.1101/2025.10.23.684201

**Authors:** Lingling Xu, Sanjeev Kumar, Kathryn M. Moore, Jacob W. Vander Velden, Faith A. Mbadugha, Meng Wang, Tiffany Hailstorks, Eric J. Sundberg, Diego E. Sastre, Mehul S. Suthar, Jens Wrammert

**Affiliations:** Division of Infectious Diseases, Department of Pediatrics, School of Medicine, Emory University, Atlanta, GA, USA; Emory Vaccine Center, Emory University School of Medicine, Atlanta, GA, USA; Centers for Childhood Infections and Vaccines, Children’s Healthcare of Atlanta, Atlanta, GA, USA; Emory Primate Center, Atlanta, GA, 30329, USA; Department of Gynecology and Obstetrics, Emory School of Medicine, Atlanta, GA, USA; Department of Biochemistry, Emory University School of Medicine, Atlanta, GA, USA

**Keywords:** Flavivirus, Zika virus, antibody-dependent enhancement, Fc receptor, FcγRI, cross-reactive antibodies, Hofbauer cells, trans-placental infection

## Abstract

Zika virus (ZIKV) can be vertically transmitted from a pregnant mother to the developing fetus, resulting in microcephaly and/or other congenital malformations. Dengue virus (DENV) cross-reactive antibodies can facilitate ZIKV placental transcytosis and enhance ZIKV infection of placenta macrophage-Hofbauer cells through binding to Fc-γ receptors (FcγRs). To understand the role of individual FcγR in antibody-mediated ZIKV placental infection, we generated a comprehensive panel of Fc-variants spanning a wide range of binding affinities to different FcγRs. We found that mutations with increased affinity to FcγRI strongly correlated with an increased frequency of infected pro-monocytic U937 and Hofbauer cells. Next, we genetically deleted individual FcγR in U937 cells, and found that the knockout of *FCGR1A* gene completely abolished ZIKV infection. In contrast, the deletion of *FCGR2B* gene showed no effect on ZIKV infection, and the deletion of *FCGR2A* gene had only a moderate impact on ZIKV infection. We further observed that FcγRI was involved in both increased ZIKV internalization and replication. Collectively, our results establish FcγRI as the key Fc receptor responsible for antibody-mediated ZIKV infection in both U937 and primary placental macrophages. These mechanistic findings not only provide insight into the importance of FcγRI in ZIKV vertical transmission but also highlight FcγRI as a potential therapeutic target, with significant implications for the development of strategies to prevent ZIKV transmission from mother to fetus.

## INTRODUCTION

Zika virus (ZIKV) is a positive-sense RNA virus that belongs to the Flaviviridae family. ZIKV is predominantly transmitted by infected Aedes mosquitoes, but can also be transmitted through sexual contact, blood transfusion, and vertical transmission from a pregnant mother to the fetus ^1^. Although ZIKV infections in adults are mostly asymptomatic or mildly symptomatic, congenital ZIKV infection is associated with an increased risk of severe neurological syndromes and congenital abnormalities, such as microcephaly ^1–3^. ZIKV was first discovered in Uganda in 1947 and was initially associated with sporadic cases of human infection until 2007 ^4^. Between 2007 and 2015, several ZIKV outbreaks were reported in Africa, Asia, the Pacific and the Americas ^1,5^. However, in 2015-2016, a ZIKV outbreak emerged in Brazil and spread rapidly to other South American countries. During this period, there was a dramatic increase in recorded cases of microcephaly and other birth defects in newborns, which was ultimately linked to ZIKV infection of the fetus during gestation ^6–10^. Given the severe outcomes associated with congenital infections, understanding the mechanisms and factors contributing to ZIKV vertical transmission is of critical importance.

ZIKV is closely related to dengue virus (DENV), another flavivirus that causes over 10 million infections each year ^11^. It has been noted that ZIKV outbreaks often occur in DENV endemic regions ^12^. The envelope (E) protein of ZIKV and DENV shares over 50% protein sequence identity and structural similarities ^13^. Numerous studies have demonstrated that DENV-specific antibodies that cross-react with ZIKV can enhance ZIKV infection in Fc gamma receptor (FcγR)-expressing cells ^14–18^. In addition, DENV-specific antibodies and DENV IgG-containing plasma have been shown to enhance ZIKV infection and disease severity in both *in vitro* and *in vivo* models ^15–17,19^.

Previously, we have shown that the ZIKV virus can cross the placental barrier using the neonatal Fc receptor (FcRn)-mediated transport system of IgG in the presence of cross-reactive antibodies in DENV-experienced individuals ^20^. Syncytiotrophoblasts, forming the outermost layer of the placenta, separating the maternal blood and the fetal compartment, are resistant to ZIKV infection^21^. However, we have previously demonstrated in a human placental explant model that cross-reactive DENV antibodies can facilitate ZIKV transcytosis across the syncytiotrophoblast layer through the FcRn ^18^. Upon accessing the placenta, ZIKV replicates primarily within cytotrophoblasts and macrophages called Hofbauer cells ^22–26^. Hofbauer cells reside in the stroma of the chorionic villi along with fibroblasts and mesenchymal cells, adjacent to the fetal vasculature^27^. Our group has previously demonstrated that cross-reactive DENV antibodies can enhance Hofbauer cell infection in an FcγR-dependent manner ^18^. Furthermore, pre-existing DENV immunity has been shown to result in enhanced ZIKV placental infection, and worsened pregnancy outcomes in both mouse and non-human primate models ^18,28–30^. These findings suggest that in DENV-experienced individuals, cross-reactive antibodies may facilitate viral seeding of the placenta and enhanced infection in Hofbauer cells, allowing ZIKV to access the fetal bloodstream.

In humans, the canonical FcγRs comprise the high-affinity receptor FcγRI (also known as CD64) and the low-affinity receptors FcγRII (FcγRIIA and FcγRIIB) (also known as CD32A and CD32B, respectively) and FcγRIII (also known as CD16) ^31–34^. FcγRI is the only receptor that binds to both monomeric IgG and immune complexes (ICs) with high affinity ^31–34^. In contrast, the low-affinity receptors only efficiently bind IgG in the form of ICs ^31–34^. The activation of FcγRs leads to a variety of effector functions, such as antibody-dependent cellular cytotoxicity (ADCC) and antibody-dependent cellular phagocytosis (ADCP) ^31–34^. Studies have shown that antibody-virus ICs can be endocytosed into FcγR-expressing cells, which results in an increased number of infected cells ^18,20,35^ and a reduction in anti-viral response ^18,36,37^. Various factors, including the type of FcγRs, IgG subclass, and antibody Fc glycosylation, determine the outcome of antibody-FcγR mediated flavivirus infection ^31,38^.

Here, we defined the key FcγRs mediating antibody-dependent ZIKV placental infection using both antibody Fc engineering and FcγR genetic modification. We found a strong positive correlation between increased ZIKV infection and antibody binding affinity to FcγRI, rather than to FcγRIIA and FcγRIIB, in both U937 and Hofbauer cells. In addition, *FCGR1A* gene knockout completely abolished ZIKV infection in U937 cells. In contrast, *FCGR2B*-knockout cells exhibited comparable ZIKV infection. *FCGR2A*-knockout cells only showed a modest decrease in ZIKV infection compared to WT cells. In addition, FcγRI enhanced ZIKV infection by facilitating both ZIKV internalization and replication. Our findings establish FcγRI as the central Fc receptor responsible for antibody-mediated ZIKV infection in both U937 and Hofbauer cells. This study advances our mechanistic understanding of antibody-mediated ZIKV pathogenesis and highlights FcγRI as a potential therapeutic target that may help guide the development of future treatment strategies for ZIKV infection in pregnant women.

## RESULTS

### FcγRI is the primary Fc receptor for antibody-mediated ZIKV infection in pro-monocytic U937 cells

To determine the contribution of each FcγR (FcγRI, FcγRII, and FcγRIII) on antibody-dependent enhanced (ADE) of ZIKV infection, we generated a panel of 13 antibody Fc mutants with a broad range of binding affinities to the different FcγRs ^39–48^ (**Figure 1 and Table S1)**. These Fc IgG1 variants, generated either by single point mutations (such as G236A, P328D, and I332E) or by multiple mutations (including AAA, EF, ADE, GASDALIE, V12, VLPLL, FTDE, LALA, LALA-PG, and GRLR), are primarily located in the lower hinge region (residues 233–239) and around the top of the CH2 domain, specifically clustering in the BC loop (residues 267–275), the FG loop (residues 324–334), and near the *N*-glycan loop (residues 292–300), involving the protein–protein interface between Fc and FcγRs (**Figure 1**). For these experiments, we used an envelope (E)-protein specific monoclonal antibody (mAb) (33.3A06), previously isolated by our group from a DENV2-infected patient ^49^, with potent neutralization activity. This antibody cross-neutralizes ZIKV ^14^, facilitates ZIKV placental transcytosis and enhances ZIKV infection of the primary tropic cell within the placenta-Hofbauer cells ^18^. As controls, we verified the expression of each variant by SDS-PAGE analysis **(Figure S1A)** and confirmed functionality of the Fab domain through analysis of the binding of each Fc variant to recombinant ZIKV E-protein by ELISA assay. All the generated Fc variants were shown to exhibit comparable binding capacity to the wild-type IgG1 construct **(Figure S1B)**.

**Figure 1.**
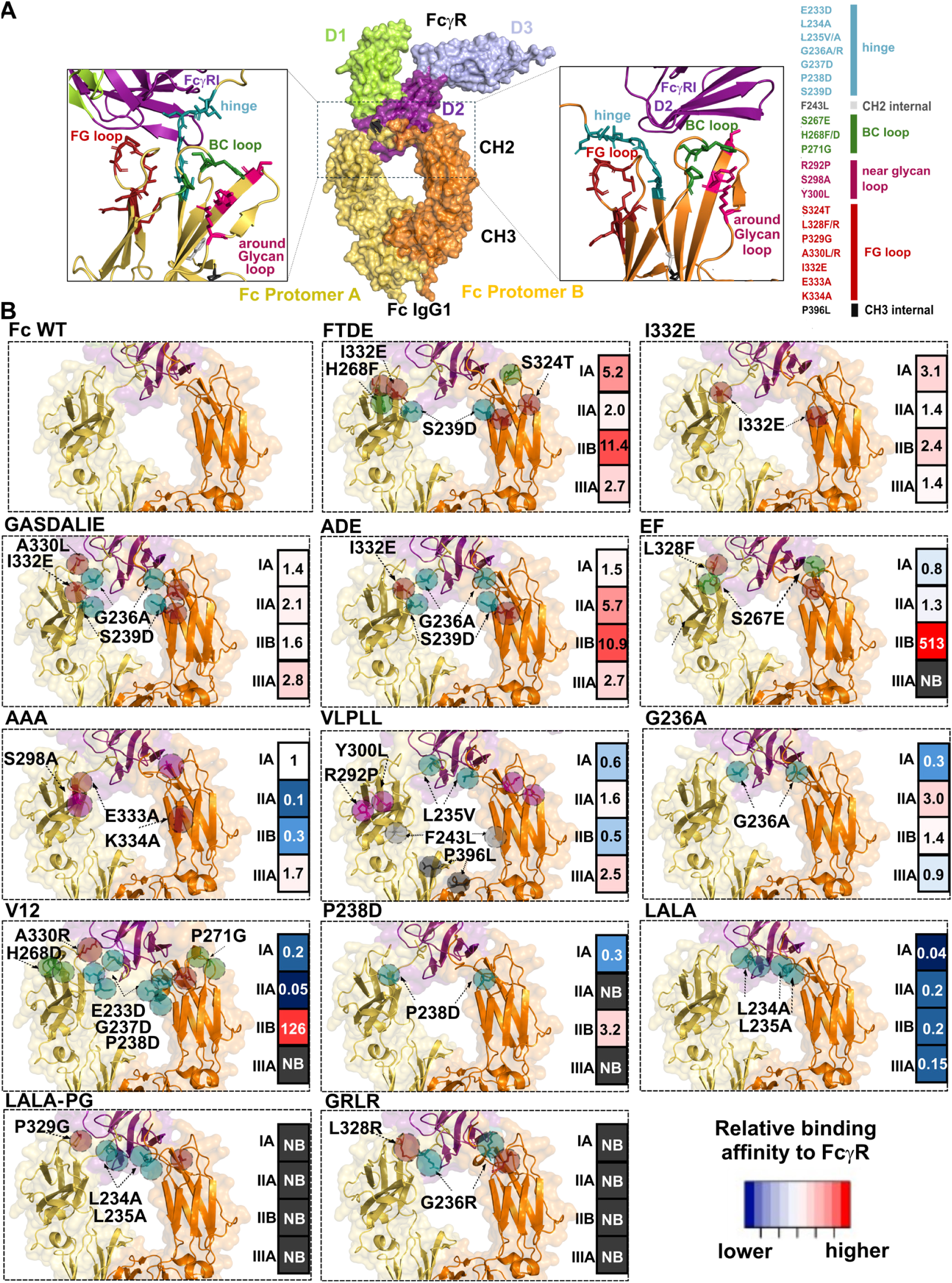
Structural representation of the 33.3A06 IgG1 Fc mutants used in the study **(A)** Surface representation of the complex Fc-FcγRI (PDB code:4W4O), indicating each region of the Fc (CH2 and CH3 domains in both protomer A (yellow) and protomer B (orange) of IgG1 Fc). It is also showing a cartoon representation of a lateral view of the crystal structure of the Fc-FcγRI complex (pdb code:4W4O), with a focus on the top of the CH2 domain of Fc from each protomer in contact with the D2 domain of FcγRI (left). A list of residues mutated in different regions of the Fc, with a color code to identify each main region that was mutagenized, is shown on the right. *N*-glycans on both Fc and FcγRI were removed to improve visibility of mutated residues. **(B)** Surface and cartoon representations of Alphafold3 models generated for each Fc variant in complex with FcγRI. The relative binding affinity of each Fc variant to each FcγR generated by BLI is indicated on the right side of each panel. Binding affinity is color-scaled: higher affinity (red), similar affinity (white) and lower affinity (blue) relative to the reference Fc WT. See also **Figure S1, Figure S2** and **Table S1**.

We then characterized the binding affinity of the 33.3A06 WT and its 13 Fc variants using Bio-Layer Interferometry (BLI) **(Table S1)** against FcγRI, FcγRIIA, FcγRIIB, and FcγRIIIA. Among the Fc variants, LALA-PG and the GRLR mutations completely abrogated binding against all four FcγRs (**Figure 1B**), which is in clear agreement with previous reports ^41,48^. The LALA mutant has been reported to have no detectable binding to FcγRs and has been extensively used in *in vivo* studies to diminish FcγRs-mediated functions. However, using BLI technique, we found that LALA showed weak but still detectable binding to all the FcγRs (Figure 1B and Table S1). The EF and V12 mutants, engineered for enhanced binding to FcγRIIB ^41,45^, showed 513-fold and 126-fold increased affinity to FcγRIIB, respectively (**Figure 1B**). The GASDALIE, G236A, and ADE mutants exhibited two- to five-fold enhanced binding affinities to FcγRIIA compared to WT IgG1 (**Figure 1B**). The FTDE mutant showed increased binding to FcγRI (5-fold), FcγRIIA (2-fold), FcγRIIB (11-fold) and FcγRIIIA (3-fold).

Next, we tested our panel of Fc variants in an *in vitro* ADE assay using the U937 monocyte cell line. We then analyzed the correlation between the antibody’s FcγRs binding affinity and ZIKV ADE. By flow cytometry, we showed that U937 cells expressed FcγRI, FcγRIIA and FcγRIIB, but not FcγRIIIA (**Figure 2A**). U937 cells are not permissive to ZIKV infection in the presence of a non-cross-reactive control mAb ^14,18^ **(Figure S4)**. However, with the addition of WT 33.3A06 IgG1, ZIKV infection increased up to 37% at 24 hours post infection (**Figure 2B**). Fc variants FTDE, I332E and EF exhibited 42%,43% and 44% peak ZIKV infection, respectively (**Figure 2B**) and a 1.3 fold increased area under curve (AUC) compared to WT with antibody concentration ranges from 0.1ng/ml-2000ng/ml (**Figure 2C**). The two variants LALA-PG and GRLR, which abolished binding to all Fc receptors (**Figure 2C, E, G)**, also showed complete abrogation of ZIKV ADE (**Figure 2B**). The LALA mutant, which is known for decreased binding to all the FcγRs ^46,47^ showed almost no infection in U937 cells (**Figure 2B**). Interestingly, we observed decreased AUC value of ZIKV ADE for all the Fc variants with reduced FcγRI binding affinity, such as VLPLL (1.6-fold), G236A (1.5-fold), V12 (2-fold), and P238D (2.3-fold) (**Figure 2B, C)**. However, the mutations that had comparable FcγRI binding affinity to WT, such as AAA, GASDALIE and ADE, exhibited equivalent ZIKV ADE compared to WT (**Figure 2B, C)**, despite their broad range of binding affinities to FcγRIIA or FcγRIIB (**Figure 2D, F, H)**. A strong correlation was observed between peak of ZIKV ADE and binding affinity to FcγRI (r=0.8392; p=0.0006), but not to FcγRIIA (r=0.3455; p=0.1496) and FcγRIIB (r=0.4406; p=0.1542) (**Figure 2E, G, I)**. These results suggest that FcγRI is the predominant receptor involved in ZIKV ADE in U937 cells.

**Figure 2.**
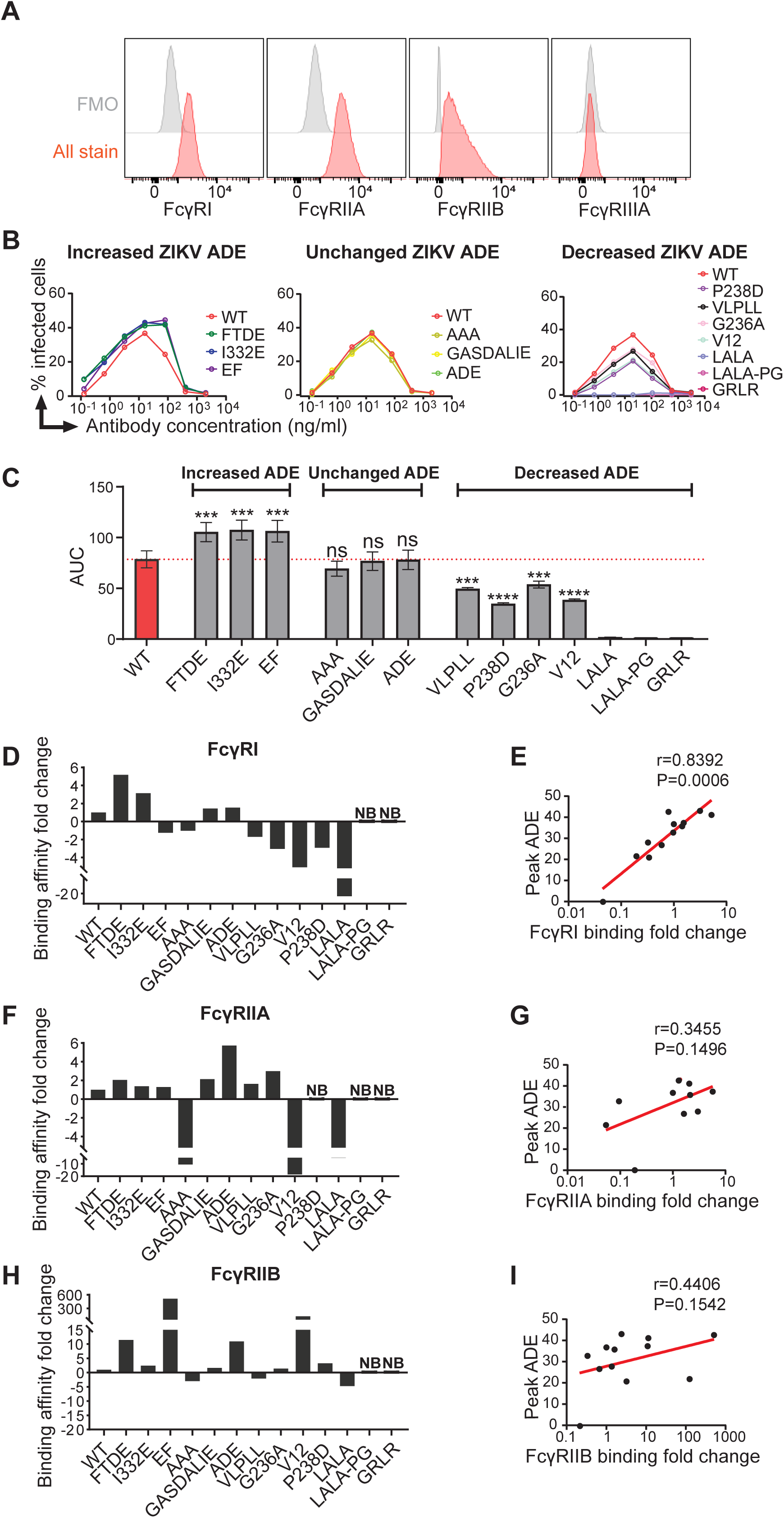
FcγRI is the primary Fc receptor for antibody-mediated ZIKV infection in U937 cells **(A)** Representative histogram showing the surface expression of FcγRI, FcγRIIA, FcγRIIB and FcγRIIIA on U937 cells as detected by flow cytometry. FMO, fluorescence minus one. **(B)** U937 cells were infected with ZIKV (MOI 0.5) and in the presence of 33.3A06 IgG1 WT and Fc variants at a range of mAb concentrations. Infected cells were detected by 4G2 staining at 24 hr post infection (hpi) and flow cytometry. Antibody dilutions were performed in singlicate. The ADE assay was repeated in three independent experiments. **(C)** Data from the same experiment as in (B), but plotted as area under the curve (AUC) to obtain better quantitative analysis. The data shown are the results of three independent experiments. Data was analyzed using 1-way ANOVA and Tukey’s multiple comparison test: ***p < 0.001, ****p < 0.0001. **(D, F, H)** Fold changes in FcγRs binding affinity of the 33.3A06 Fc variants. K_D_ values of the WT IgG1 and Fc variants binding to recombinant human FcγRI, FcγRIIA and FcγRIIB were determined by BLI. Relative binding affinity=K_D_ (WT)/K_D_ (variant). Variants with negative values show decreased binding affinity. NB, no detectable binding. **(E, G, I)** Correlations between peak ZIKV ADE and FcγRI, FcγRIIA and FcγRIIB relative binding affinity of the 33.3A06 IgG1 WT and Fc variants. Correlations were analyzed by Spearman’s Rho on log-transformed data. P-value was considered significant if <0.05. See also **Figure S3, S4 and S5.**

### *FCGR1A* knockout abrogates ZIKV ADE in U937 cells

We found that U937 cells and Hofbauer cells express FcγRI, FcγRIIA and FcγRIIB, and all of which can bind to immune complexes ^31,50^. To further dissect the contribution of each FcγRs on antibody-dependent ZIKV infection, we generated U937 cells knock-outs (KO) for *FCGR1A*, *FCGR2A* and *FCGR2B* using a CRISPR-Cas9 strategy. Briefly, stable *FCGR* KO cell lines were generated by transducing sgRNA containing lentiviral particles, followed by single-cell sorting of GFP+ cells. The KO status of the clones was validated by flow cytometry. We were able to specifically ablate expression of the individual FcγRs without affecting the expression of the other Fc receptors (**Figure 3A**). Next, we conducted *in vitro* ZIKV ADE assays using either WT or FcγR knockout cell lines and the WT 33.3A06 IgG1 mAb. Interestingly, with mAb concentrations ranging from 0.005 ng/ml-10ug/ml, *FCGR1A* knockout cells showed completely abolished ZIKV infection (**Figure 3B**). The *FCGR2A* knockout cells showed a decrease in peak ADE (2-fold) and AUC (1.8-fold) compared to WT cells (**Figure 3B, C)**. In contrast, *FCGR2B* knockout cells exhibited comparable peak ZIKV ADE and AUC to those of WT cells (**Figure 3B, C)**.

**Figure 3.**
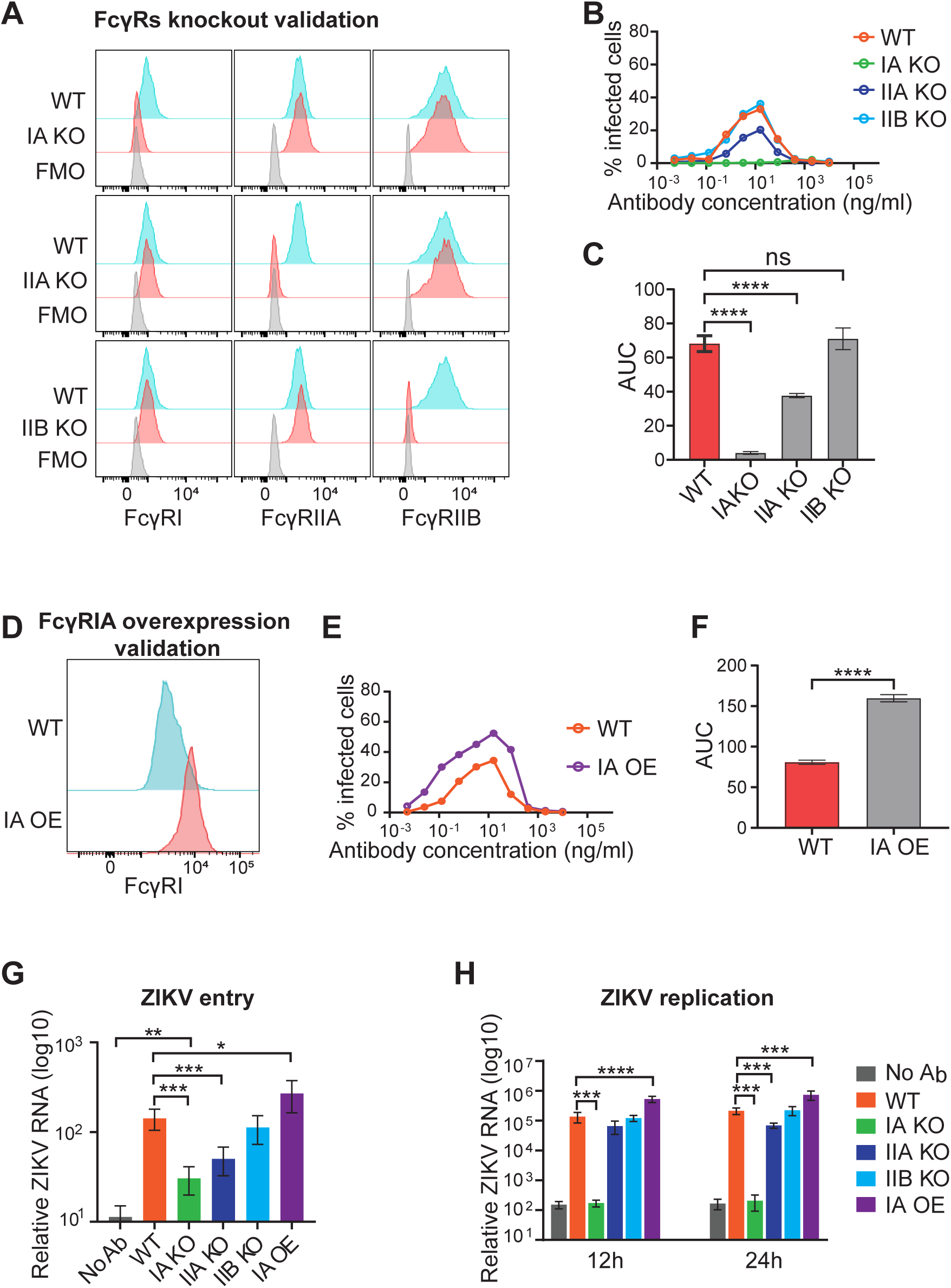
FcγRI-mediated enhanced ZIKV infection in U937 cells through both pre- and post-entry mechanisms. **(A)** Representative histogram showing the surface expression of FcγRI, FcγRIIA and FcγRIIB in WT, *FCGR1A* knockout, *FCGR2A* knockout and *FCGR2B* knockout U937 cell lines, respectively, as detected by flow cytometry. **(B)** WT, *FCGR1A* knockout, *FCGR2A* knockout and *FCGR2B* knockout U937 cell lines were infected with ZIKV (MOI 0.5) in the presence of 33.3A06 IgG1 mAb. Infected cells were detected by 4G2 staining and flow cytometry at 24 hpi. Data shown are representative of three independently performed experiments. **(C)** Data from the same experiment as in (B), but plotted as AUC to obtain better quantitative analysis. The data shown are the results of three independent experiments. Data was analyzed using 1-way ANOVA and Tukey’s multiple comparison test: ****p < 0.0001. **(D)** Representative histogram showing surface expression of FcγRI on WT and FcγRI overexpressed U937 cell lines as detected by flow cytometry. **(E)** WT and FcγRI overexpressed U937 cell lines were infected with ZIKV (MOI 0.5) in the presence of 33.3A06 IgG1 mAb at a range of mAb concentrations. Infected cells were detected by 4G2 staining and flow cytometry at 24 hpi. Data shown are representative of three independently performed experiments. **(F)** Data from the same experiment as in (E), but plotted as AUC to obtain better quantitative analysis. The data shown are the results of three independent experiments. Data were analyzed by Student’s t-test: ****p < 0.0001. **(G)** U937 cells were infected with ZIKV (MOI 1) alone or in the presence of 33.3A06 IgG1 mAb at 16ng/ml. ZIKV RNA from internalized virus was measured by qRT-PCR. **(H)** ZIKV RNA at 12hpi and 24hpi was measured by qRT-PCR. Data are representative of three independently performed experiments. Data were analyzed by Student’s t-test: *p<0.05, **p < 0.01, ***p < 0.001, ****p < 0.0001. Data are presented as mean values ± SD. IA OE, FcγRI overexpression; WT, wild type; IA KO, *FCGR1A* knockout; IIA KO, *FCGR2A* knockout; IIB KO, *FCGR2B* knockout.

To determine how antibody factors such as epitope specificity, neutralizing potency and virion binding affinity impact antibody-mediated ZIKV infection, we also evaluated the ADE effect of a panel of cross-reactive mAbs **(Figure S5A)**, which vary in neutralization, epitope specificity ^14,49,51,52^ and ZIKV binding affinity **(Figure S5B)**. Using this panel of mAbs, we conducted *in vitro* ZIKV ADE assays using WT and *FCGR* knockout cell lines. All the cross-reactive antibodies enhanced ZIKV infection in WT U937 cells, but with varied peak infection or enhancement antibody concentrations. Like 33.3A06, we did not observe successful ZIKV infection in *FCGR1A* knockout cells for all the tested mAbs **(Figure S5C)**. *FCGR2B* knockout cells showed comparable ZIKV ADE to WT cells across the tested mAbs **(Figure S5C)**. The *FCGR2A* knockout cells showed an approximately 2-fold decreased peak ADE compared to WT cells for all the test mAbs, except for the E-protein domain III (EDIII)-specific antibody Z004, which was comparable to WT.

To assess the effect of over-expression of FcγRI on ZIKV ADE, we cloned the FcγRIA alpha chain and *FCER1G* genes linked by the nucleotides of the P2A self-cleaving peptide into a lentiviral vector. A stable FcγRI overexpressing cell line was generated by transducing packaged lentiviral particles into WT U937 cells (**Figure 3D**). The *in vitro* ZIKV ADE assay using the FcγRI overexpressing cell line exhibited a more potent ZIKV infection, with the peak infection increasing from 34% to 52% (**Figure 3E**) and a 2-fold increase in AUC (**Figure 3F**), compared to WT cells. Altogether, these results indicate that FcγRI is the primary Fc receptor responsible for ZIKV ADE in U937 cells, while FcγRIIA may have a minor role in this process.

### Both pre- and post-internalization mechanisms impact FcγRI-mediated enhanced ZIKV infection in U937 cells

To understand the mechanism/s by which FcγRI contributes to antibody-mediated enhanced ZIKV infection, we performed a viral internalization assay using WT, *FCGR* knockout cells and FcγRI overexpressing cells with ZIKV alone or in the presence of the 33.3A06 IgG1 mAb by measuring viral RNA at 2 hours post virus entry. As expected, we observed significantly increased viral internalization in ZIKV-infected WT cells compared with virus alone. We also observed around 80% decreased viral internalization in *FCGR1A* knockout cells (**Figure 3G**) and around 65% decreased viral internalization in *FCGR2A* knockout cells compared to WT cells (**Figure 3G**). In addition, we observed around two-fold increased viral internalization in FcγRI-overexpressing cells compared with WT cells (**Figure 3G**). *FCGR2B* knockout cells showed comparable viral internalization to WT cells (**Figure 3G**). To determine whether FcγRs impact ZIKV replication after internalization into cells, we next quantified ZIKV viral RNA at 12 hours and 24 hours after entry. Extensive ZIKV replication was observed in WT cells, *FCGR2A* knockout cells, *FCGR2B* knockout cells and FcγRI overexpressing cells (**Figure 3H**). Despite a higher level of viral internalization than without antibody, *FCGR1A* knockout cells showed a similar level of viral RNA to the virus alone group at both 12h and 24h (**Figure 3G and H**), illustrating that viral replication in the cells was significantly impacted by the lack of FcγRI. In addition, we observed around 4-fold increased viral RNA in FcγRI-overexpressing cells compared to WT cells at 12h and 24h post entry (**Figure 3H**). We also observed a 3-fold decrease in viral RNA in the *FCGR2A* knockout cells compared to WT cells at 24h (**Figure 3H**). These results suggest that FcγRI not only promotes ZIKV virus internalization but is also critical for ZIKV replication in the cells.

### FcγRI is the primary Fc receptor for antibody-mediated ZIKV infection in placental Hofbauer cells

Evidence from both *in vitro* and *in vivo* studies indicates that DENV cross-reactive antibodies enhance ZIKV infections during pregnancy, and antibody-mediated ZIKV infection is a critical mechanism of ZIKV vertical transmission ^18,28–30^. Next, we sought to determine which FcγRs impact ZIKV ADE in Hofbauer cells isolated from full-term placentas. Like U937 cells, Hofbauer cells also express FcγRI, FcγRIIA and FcγRIIB, but with relatively higher FcγRI expression compared to U937 cells (**Figure 4A and S6)**. We evaluated the effects of the panel of Fc mutants described above on ZIKV infection in primary Hofbauer cells. We used a constant antibody concentration of 16 ng/ml due to the limited quantity of primary cells that could be obtained from one placental tissue. In contrast to the U937 cells, around 2% of primary Hofbauer cells were infected by ZIKV at an MOI of 0.25, even in the absence of mAbs (**Figure 4B**). The WT 33.3A06 IgG1 caused enhanced ZIKV infection to around 33% (**Figure 4B**). The LALA-PG and GRLR mutants showed a comparable baseline infection rate to the no antibody group (Figure 4B). Similar to U937 cells, the three variants with unchanged affinity for FcγRI, ie. AAA, GASDALIE and ADE, caused comparable ZIKV ADE to WT IgG1 in Hofbauer cells. The mutations with decreased FcγRI binding affinity, ie. G236A, V12, and P238D caused 24.5%, 23.5% and 26.3% Hofbauer cell infection, respectively (**Figure 4B**). The mutant VLPLL exhibited a 1.7-fold decreased affinity for FcγRI compared to WT IgG1, but comparable ZIKV infection to WT IgG1 in Hofbauer cells. Interestingly, the LALA mutation, which completely ablated ZIKV infection in U937 cells, enhanced ZIKV infection in Hofbauer cells to around 11% (**Figure 4B**). The different ZIKV infection outcomes in U937 and Hofbauer cells caused by LALA and VLPLL are possibly due to a higher level of FcγRI expression in Hofbauer cells, which makes them less sensitive to decreased

**Figure 4.**
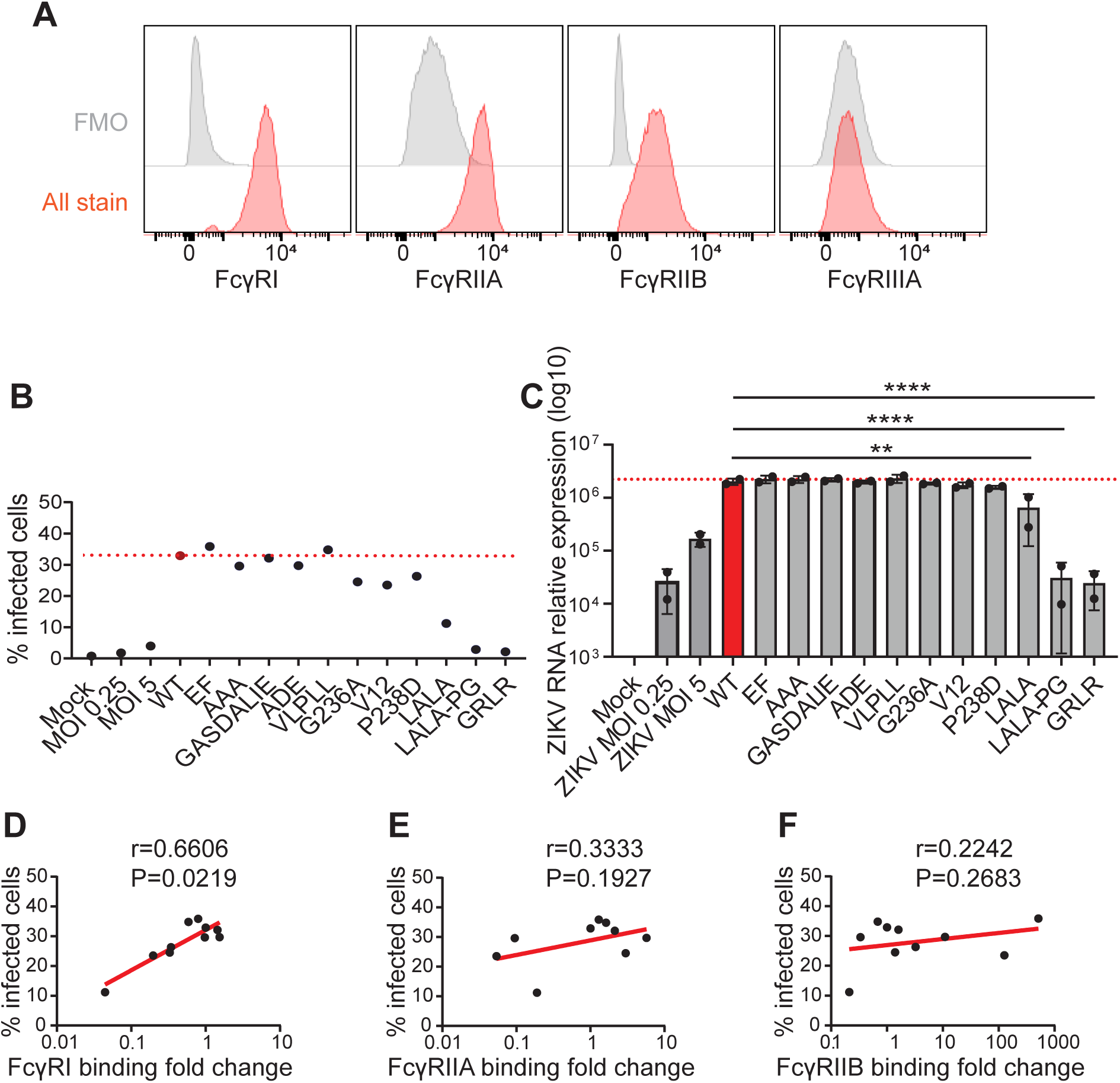
FcγRI is a main receptor for ZIKV ADE in Hofbauer cells. **(A)** Representative histogram showing the surface expression of FcγRI, FcγRIIA, FcγRIIB and FcγRIIIA on Hofbauer cells as detected by flow cytometry. FMO, fluorescence minus one. **(B)** Hofbauer cells were infected with ZIKV (MOI 0, 0.25 or 5) alone or MOI 0.25 in the presence of mAbs at 16ng/ml. Infected cells were detected by 4G2 staining and flow cytometry at 24 hpi (biological replicates ± SD). Representative experiment from n = 2 donors. **(C)** Hofbauer cells were infected with ZIKV (MOI 0, 0.25 or 5) alone or MOI 0.25 in the presence of mAbs at 16ng/ml. ZIKV RNA was measured by qRT-PCR at 24 hpi (biological replicates ± SD). Representative experiment from n = 2 donors. Data was analyzed using 1-way ANOVA and Tukey’s multiple comparison test: *p<0.05, **p < 0.01, ****p < 0.0001. **(D, E, F)** Correlations between ZIKV infection in Hofbauer cells and FcγRI, FcγRIIA and FcγRIIB relative binding affinity of the 33.3A06 IgG1 and Fc variants. Correlations were analyzed by Spearman’s Rho on log-transformed data. P-value was considered significant if <0.05. See also **Figure S4 and S5.**

FcγRI binding affinity compared to U937 cells. We also measured ZIKV RNA levels by RT-PCR and found that LALA, LALA-PG and GRLR mutants caused a significant decrease in ZIKV RNA in Hofbauer cells (**Figure 4C**). Finally, we observed a strong correlation (r=0.6606; p=0.0219) between FcγRI binding affinity and ZIKV ADE (**Figure 4D, E, F)**. Taken together, these results suggest that FcγRI is the predominant receptor of ZIKV ADE not only in the U937 cell line, but also in primary placental Hofbauer cells.

## DISCUSSION

The impact of pre-existing immunity in shaping the clinical severity of flavivirus infections, such as ZIKV, remains unclear. Prior studies suggest that cross-reactive antibodies can mediate vertical transmission of ZIKV across the placental barrier and subsequent infection of placental macrophages in an FcγR-dependent manner ^18^. Virus-specific antibodies facilitate increased viral entry and replication in host cells through antibody-dependent enhancement (ADE) ^53–55^. The precise Fcγ receptors mediating this process in human immune and placental cells remain poorly defined. Monocytes and macrophages are the primary targets of antibody-mediated ZIKV and DENV infection^56,57^. These cells express FcγRI, FcγRIIA, FcγRIIB and FcγRIIIA ^31,56^. Previous reports have shown that blocking FcγRI and FcγRIIA with monoclonal antibodies significantly reduce antibody-mediated DENV or ZIKV infection in FcγRs-bearing cell lines ^56–60^. However, there have been conflicting results in terms of which FcγR is the primary driver of antibody-mediated DENV and ZIKV infection. This might be due to incomplete blocking or poorly specific activity of the blocking antibodies ^37,57,61^.

To overcome the limitation of blocking antibodies, we approached this lack of understanding using genetic ablation and antibody Fc engineering approaches. To determine which receptor(s) mediate ZIKV infection in primary Hofbauer cells and the U937 cell line, we generated a panel of engineered IgG1 Fc mutants with a broad range of FcγR binding affinities, using the potent DENV and ZIKV cross-neutralizing human antibody 33.3A06 ^14,49^. This panel of mutant antibodies and their binding to different FcγRs have not been comprehensively characterized in a side-by-side fashion previously. Here, we characterized the binding affinity and functional characteristics of the Fc mutant panel of the 33.6A06 mAb on antibody-mediated ZIKV infection using primary Hofbauer cells and the U937 cell line. We observed a strong positive correlation between antibody affinity for FcγRI and ZIKV infection in both U937 and Hofbauer cells, while FcγRIIA and FcγRIIB did not correlate with ZIKV infection. Thus, mutants with enhanced FcγRIIA binding, such as GASDALIE and ADE, did not alter ZIKV infection. Similarly, mutations enhancing FcγRIIB binding, such as the EF mutation, showed comparable ZIKV infection to that of WT IgG1 in Hofbauer cells. These results support a major role for FcγRI during ZIKV placental infection.

Primary Hofbauer cells cannot be genetically modified, due to their relatively short lifespan *in vitro*. In light of this, the U937 monocytic cell line is a useful model for studying antibody-mediated ZIKV infection *in vitro*, as it can be readily modified genetically. However, it is worth pointing out that the two cell types exhibit different responses to ZIKV infection in a few key aspects. Hofbauer cells can be directly infected by ZIKV in the absence of antibodies, due to the expression of ZIKV entry receptors such as DC-SIGN ^62,63^. In contrast, U937 cells are completely resistant to ZIKV infection in the absence of antibodies, making this cell line a good model for studying antibody-mediated flavivirus infection. In addition, the LALA mutant does not trigger ZIKV infection in U937 cells, but does enhance ZIKV infection modestly in Hofbauer cells. This difference could be due to the different receptor density on these cells, allowing Hofbauer cells to become infected even through the low affinity FcγR-IgG1 LALA interaction. These results suggest that future passive immunotherapy for ZIKV infection in pregnancy should consider GRLR or LALA-PG mutations, rather than LALA, to eliminate potential antibody-mediated infection.

To further define the involvement of Fcγ receptors in antibody mediated ZIKV infection, we developed knock-out lines of U937 cells, using CRISPR-Cas9 approaches. Using these novel lines, we could show that ZIKV was unable to infect the U937 cells in the absence of *FCGR1A*. Deletion of *FCGR2B* had no impact on the infection rate of U937 cells, while *FCGR2A* showed a minor impact on the magnitude of infection. In addition, by carefully measuring entry and replication kinetics in these knockout lines, we could show that the FcγRI receptor is critical for successful ZIKV entry in U937 cells. We also observed a somewhat reduced ZIKV internalization in *FCGR2A* knockout cells, matching the results above, and also matching previous data showing that FcγRIIA can impact DENV infection of U937 and K562 cells ^59,64,65^. Mechanistic experiments are underway to address exactly how the FcγRI interaction modulates viral replication. Taken together, these findings show that FcγRI is the dominant receptor for antibody-mediated ZIKV infection in U937 and Hofbauer cells.

In summary, we show that FcγRI is the predominant Fc receptor for antibody-mediated ZIKV infection in both U937 cells and primary human Hofbauer cells. We demonstrated that FcγRI promoted antibody-mediated ZIKV infection by enhancing both virus entry and replication. These findings highlight the critical role FcγRI may have in ZIKV vertical transmission. Our results suggest that therapeutic strategies aimed at blocking FcγRI function may be more effective in mitigating ZIKV pathogenesis. Future studies will investigate the molecular signaling pathway involved in FcγRIA-mediated virus replication. These findings advance our understanding of ZIKV-host interactions and open new avenues for targeted interventions to prevent ADE-related complications during ZIKV infection, particularly in pregnant women.

## LIMITATIONS OF THE STUDY

One limitation of this study is the relatively small number of placental samples used for testing the effect of 33.3A06 Fc mutants on ZIKV placental infection. It is possible that inter-individual variability may not have been fully captured among the donors analyzed. Further, primary Hofbauer cells are difficult to purify in large numbers and cannot be genetically modified due to their relatively short life span *in vitro*. Due to this limitation, we only tested a single antibody concentration (16ng/ml) for Hofbauer cell ADE assays and used monocyte cell line U937 cells to study the mechanism of FcγRI-mediated ZIKV infection in more detail.

## ACKNOWLEDGEMENTS

We thank the Emory Chemical Biology Discovery Center for providing access and support in conducting the BLI assays. We also thank Devyani Joshi for critical review of the manuscript. This work was funded in part by the NIH grant R01 AI149486 (J.W and M.S.S) and 5U19AI057266-13REVIS Supplement (M.S.S and J.W), by the Pediatric Research Alliance Center for Childhood Infections and Vaccines and Children’s Healthcare of Atlanta. We thank DBT/Wellcome Trust India Alliance Early Career Fellowship grant IA/E/18/1/504307 awarded to S.K.

## AUTHOR CONTRIBUTIONS

Experimental work, data acquisition, data analysis and figure generation by L.X., S.K., K.M., J. V., F.M. M.W., D.S., M.S.S., J.W. Conceptualization by L.X., S.K., K.M., M.S.S and J.W. Manuscript writing by L.X., D.S and S.K. Manuscript review & editing by L.X., S.K., K.M., D.S., M.S.S and J.W. Funding Acquisition by M.S.S and J.W. All authors have read and accepted the manuscript.

## DECLARATION OF INTERESTS

The authors declare no competing interests.

## MATERIALS AND METHODS

### EXPERIMENTAL MODEL AND STUDY PARTICIPANT DETAILS

#### Human subjects

Human term placentae (>37 weeks gestation) were collected from hepatitis B, HIV-1 seronegative women (>18 years of age) immediately after elective cesarean section without labor from Emory Midtown Hospital, Atlanta, GA. This study was approved by the Emory University Institutional Review Board (IRB 000217715). Written informed consent was obtained from all donors before cesarean section and sample collection. Placenta samples were de-identified before being transferred to laboratory personnel for primary Hofbauer cell isolation.

#### Primary Hofbauer cell and monocyte isolation

Hofbauer cells were isolated from membrane-free villous full-term placenta and purified using anti-CD14 magnetic beads as previously described ^66^. After isolation, Hofbauer cells were cultured in complete RPMI medium consisting of 1x RPMI (Corning Cellgro), 10% FBS (Optima, Atlanta Biologics), 2mM L-glutamine (Corning Cellgro), 1mM sodium pyruvate (Corning Cellgro), 1x Non-essential AminoAcids (Corning Cellgro), 1x antibiotics (penicillin, streptomycin, amphotericin B; Corning Cellgro) at 37°C and 5% CO2.

#### ZIKV virus

ZIKV strain PRVABC59 was obtained from the Centers for Disease Control and Prevention (CDC) and was passaged by infecting Vero cells (ATCC; CRL-1586) at a multiplicity of infection (MOI) of 0.1 in serum-free MEM (Life Technologies Gibco) as previously described ^14,18^. Virus was titered by focus forming assay on Vero cells as previously described and stored at −80 °C ^23^.

#### Cell lines

U937 cells (ATCC; CRL-1593.2) were maintained in RPMI-1640 supplemented with 10% fetal bovine serum (FBS), 2mM L-Glutamine (Corning), 1mM HEPES (Corning), 1mM sodium pyruvate (Corning), 1X Non-essential Amino Acids (NEAA) (Corning) and 1X penicillin/streptomycin (Corning). HEK-293T (ATCC; CRL-3216) cells were cultured in standard medium [Dulbecco’s modified Eagle’s medium (DMEM) supplemented with 10% FBS, 2mM L-Glutamine, and 1X penicillin/streptomycin. Vero cells (ATCC; CRL-1586) were cultured in MEM medium supplemented with 10% FBS, 2mM L-Glutamine, 1X NEAA, 1mM HEPES, 1mM sodium pyruvate and 1X penicillin/streptomycin. All the above cells were cultured at 37°C with 5% CO_2_.

## METHOD DETAILS

### Generation of Fc mutant antibodies

The mAb 33.3A06 IgG1 plasmids were generated by cloning the 33.3A06 V_H_ and V_L_ genes into the human IgG1 heavy chain expression vector and human lambda light chain expression vector, respectively, as previously described ^49,67^. The 33.3A06 IgG1 Fc mutants, including H268F/S324T/ S239D/I332E (FTDE) ^39^, I332E ^40^, S267E/L328F (EF) ^41^, S298A/E333A/K334A (AAA) ^42^, G236A/S239D/A330L/I332E (GASDALIE) ^43^, G236A/S239D/I332E (ADE) ^40^, L235V/F243L/R292P/Y300L/P396L (VLPLL) ^44^, G236A ^40^, E233D/G237D/P238D/H268D/P271G/A330R (V12) ^45^, P238D ^45^, L234A/L235A (LALA) ^46,47^, L234A/L235A/P329G (LALA-PG) ^48^, G236R/L328R (GRLR) ^41^, were generated by replacing the constant region of the wild-type IgG1 heavy chain expression vector with synthesized constructs (Twist Bioscience) containing the corresponding mutations (see Figure 1A). The mutants were produced as described below.

### Structural modeling of Fc variants in complex with Fcγ receptors

Protein structure models were predicted using AlphaFold 3 (https://doi.org/10.1038/s41586-024-07487-w) through the Alphafold server. Protein sequences of each Fc IgG1 variant and ectodomains of human FcγRs (Uniprot codes: P12314 for FCGR1, P12318 for FCG2A, P31994 for FCG2B, P08637 for FCG3A) in FASTA format were submitted as input, and the models were generated using default settings without template restrictions. The top-ranked models according to the AlphaFold internal confidence score (pLDDT) were selected for downstream analyses. Model visualization and structural alignments were performed using PyMOL v3.1.

### Monoclonal Antibody Production

The human monoclonal antibodies (mAbs) used in these experiments were generated as previously described ^49^. Briefly, the variable heavy and light chain sequences encoding IgG1 expression plasmids were transiently expressed in expi293F cells and secreted IgG antibodies were purified from supernatants using protein A coupled sepharose beads. The variable heavy and light chain sequences of control antibodies EDE1 C10 ^51^ and Z004 ^52^ were synthesized (Twist Bioscience) and cloned into human IgG1 expression vectors based on PDB IDs 4UT9 and 6UTA respectively. The Pan-flavivirus anti-envelope mAb 4G2 was isolated from the supernatant of mouse hybridoma (D1-4G2-4–15; ATCC HB-112) grown in hybridoma medium (Thermofisher Scientific, 12040077), and purified using protein G column (GE Life Science) according to the manufacturer’s recommendations. The antibodies were stored in 1x PBS for in-vitro cell culture assays and with 0.05% sodium azide at 4°C for the binding assays.

### ELISA

The binding potential of mAbs to ZIKV E protein was determined by ELISA as previously described ^14,49^. Briefly, the recombinant ZIKV E protein (Memphis; R01635) was coated on Nunc Maxisorp ELISA plates (eBioscience; 44-2404) at a concentration of 1ug/ml in PBS overnight at 4 °C. The plates were washed extensively with PBS containing 0.05% Tween-20 (PBS-T) and blocked with PBS-T containing 10% FBS (PBS-T-FBS) for 1.5 h. mAbs with Serial 3-fold dilution in PBS-T-FBS were added to the plates and incubated at room temperature for 1.5 h. The ZIKV E protein specific IgG signals were detected by incubating with peroxidase-conjugated anti-human IgG (Jackson ImmunoResearch; 109-036-098) for 1 h. The plates were washed and developed using an o-phenylenediamine substrate (Sigma; P8787) in 0.05M phosphate-citrate buffer (Sigma Aldrich; P4809) containing 0.012% hydrogen peroxide. The absorbance values were measured at OD490. The absorbance values were plotted against the concentration of antibodies. The whole virus capture ELISA was performed as previously described ^14^. Briefly, the plates were coated with the pan-flavivirus 4G2 mAb overnight at 4 °C. After blocking, ZIKV was added for 1h. Plates were washed and serially diluted mAbs were added for 1.5 h, followed by the addition of peroxidase-conjugated anti-human IgG and development using an o-phenylenediamine substrate. The absorbance values were detected at 490 nm and the minimum effective concentration for binding was determined as the concentration required to obtain three times the signal obtained with an irrelevant mAb.

### Measurement of antibody-FcγRs binding kinetics by biolayer interferometry (BLI)

Octet BLI was performed using an Octet Red384 instrument (ForteBio Inc.). For measuring the binding affinity of human IgG1 Fc variants and human FcγRs, soluble human FcγRI (Thermofisher; A42518), FcγRIIA (Thermofisher; A42527), FcγRIIB (Thermofisher; A42520) and FcγRIIIA (Thermofisher; A42536), diluted at 2 μg/ml in 10 mM sodium acetate, pH 5 or 6 (pH 6 for human FcγRI and FcγRIIB; pH 5 for human FcγRIIA and FcγRIIIA) were immobilized on AR2G sensors (Sartorius; 18-5092) using AR2G Reagent Kit (Sartorius; 18-5095). The binding kinetics were measured with serial two-fold dilution of mAbs in 1X kinetics buffer (Sartorius;18-1105). Association time was 300 s followed by a 600 s dissociation step by immersing the sensors in kinetics buffers. Background binding was subtracted using reference controls and affinity constants (K_D_) were calculated based on their global fit to a 1:1 Langmuir binding model using Octet data analysis software version 12.0.

### Antibody Dependent Enhancement assay

Antibody dependent enhancement (ADE) assay was performed as previously described ^14,49^. Briefly, serially diluted mAbs were incubated with ZIKV with an MOI of 0.5 for 1 h at 37 °C. The immune complexes were added in a 96-well plate containing 2×10^4^ U937 cells (ATCC; CRL-1593.2) per well in RPMI-1640 with 10% (vol/vol) FBS. The Cells were infected for 24 h at 37 °C. The cells were then washed and fixed/permeabilized using BD intracellular staining reagents Fix/Perm Solution (BD; 51–2090KZ) and Perm/Wash Buffer (BD; 51–2091KZ) according to the manufacturer’s protocol. Cells were stained with 4G2 antibody for 1h followed by a secondary anti-mouse IgG AF488 (Life Technologies; A11029) for 25min. The cells were washed twice with Perm/Wash buffer and once with 1x PBS, and then re-suspended in 1x PBS. The percentage of infected (4G2^+^) cells was determined using flow cytometry using the Cytek Aurora flow cytometer followed by analysis using FlowJo software v10.

### ZIKV infection of primary cells

Hofbauer cells were infected immediately following isolation as previously described ^18^. Briefly, Monoclonal antibodies (mAb) were diluted in 1x PBS to the desired concentrations and mixed 1:1 with ZIKV at MOI of 0.25. mAb:ZIKV immune complexes were incubated at 37 °C for 1 hour. Cells were then infected with 200ul mAb:ZIKV complexes, or with ZIKV alone at MOI of 0.25 or 1 at 37°C for 1 hour. Hofbauer cells were washed once with warm RPMI to remove residual immune complexes and resuspended in complete RPMI medium. The cells were infected for 24 h at 37 °C prior to subsequent analysis.

### Quantitative Real Time-PCR

RNA quantification using RT-PCR was conducted as previously described ^18^. Briefly, RNA was isolated from mock-or ZIKV-infected Hofbauer cells or U937 cells using the Quick-RNA MiniPrep Kit (Zymo Research) by the manufacturer’s instructions. Purified RNA was reverse transcribed using random primers with the High-Capacity cDNA Reverse Transcription Kit (Applied Biosystems). Hofbauer cells gene expression and ZIKV viral RNA were quantified by qRT-PCR TaqMan Gene Expression Master Mix (Applied Biosystems) by the manufacturer’s instructions using ZIKV-specific primers and probe set. Host gene expression and ZIKV RNA in U937 cells were performed using SYBR green. ZIKV E gene probe sequences: ZIKV1107 (FAM AGCCTACCTTGACAAGCAATCAGACACTCAA-TAMRA), ZIKV 1086 (5-CCGCTGCCCAACACAAG-3’) and ZIKV 1162c (5’-CCACTAACGTTCTTTTGCAGACAT-3’). All primers were purchased from Integrated DNA Technologies (IDT). Primer sequences used to identify expression of host genes (forward and reverse, 5’---3’): GAPDH: GGAGCGAGATCCCTCCAAAAT and GGCTGTTGTCATACTTCTCATGG; CT values were normalized to the reference gene GAPDH and represented as fold change over values from time-matched mock samples using the formula 2−ΔΔCT. The qRT-PCR experiments were performed on an Applied Biosystems 7500 Real-Time PCR System.

### Flow Cytometry

Hofbauer cells staining were conducted as described previously ^18^. Briefly, the cells were stained for surface markers for 20min on ice using the following anti-human antibodies in FACS buffer: CD14 (M5E2; Biolegend), FcγRI (10.1; Biolegend). FcγRIIA (IV.3; Stem cell technologies),FcγRIIB (2B6; Creative Biolabs) and FcγRIIIA (3G8; Biolegend). Cells were washed with PBS and stained with Ghost 780 viability dye (TONBO Biosciences) for 20min on ice. Cells were washed with PBS and fixed with 1x Transcription Factor Fix/Perm (TONBO Biosciences) for 20min on ice and permeabilized by washing twice with 1x Flow Cytometry Perm Buffer (TONBO Biosciences). To perform intracellular staining of ZIKV E protein, cells were re-blocked for 5min on ice with Human TruStain FcX in Perm Buffer and stained with 4G2-APC antibody in Perm Buffer for 20min on ice. Cells were washed twice with 1x Flow Cytometry Perm Buffer and once with PBS. Flow cytometry samples were re-suspended in 1x FACS and run on an LSR-II flow cytometry machine and analyzed using FlowJo version 10.

U937 cells were stained for surface markers for 30 min at 4 °C using the following anti-human antibodies in FACS buffer: FcγRI (10.1; Biolegend), FcγRIIA (IV.3; Stem cell technologies), FcγRIIB (2B6; Creative Biolabs) and FcγRIIIA (3G8; Biolegend). Cells were washed with 1x PBS and stained with LIVE/DEAD fixable Yellow viability dye (Thermo Fisher Scientific). The cells were washed with 1x FACS buffer and fixed with 2% paraformaldehyde at 4 °C for 10min. The cells were then washed and re-suspended with 1x FACS buffer. Data were acquired on a BD FACSymphony A5 and analyzed using FlowJo version 10.

### Generation of *FCGR* gene knockout cells

A CRISPR/Cas9 approach was used to target the genes of FcγRIA, FcγRIIA or FcγRIIB in the U937 cell line^68^. Briefly, the CRISPR sgRNA All-in-One Lentiviral Vector: pLenti-U6-sgRNA-SFFV-Cas9-2A-GFP (Applied Biological Materials Inc; C442), pMD2.G (Addgene; 12259) and pxPAX2 (Addgene; 12260) were co-transfected (7.5ug lenti-vector, 3ug pMD2.G and 4.5ug pxPAX2) into HEK293T cells using PEI-Max transfection reagent (1 mg/ml; Polysciences) to generate lentivirus. The sgRNA sequences targeting *FCGR1A*, *FCGR2A* and *FCGR2B* were: CTGGGAGCAGCTCTACACAG, TGCTGAAACTTGAGCCCCCG and TGGAGCACGTTGATCCACTG respectively. A scrambled sgRNA sequence without targeting any gene served as a control. To generate knockout cell lines, U937 cells were transduced with the generated lentivirus and incubated at 37 °C for 72 h. Single GFP^+^ cells were sorted into 96-well plates and incubated at 37 °C for 2 weeks. Successful *FCGR1A, FCGR2A and FCGR2B* knockout cell lines were validated using flow cytometry.

### Generation of FcγRI overexpressing U937 cells

The genes of FcγRI alpha chain (NM_000566.3) and FCER1G (NM_004106.2) were linked by the nucleotides of the P2A self-cleaving peptide and cloned into a lentiviral vector: pLenti-puro (Addgen, 39481). The lentivirus was produced as described above. The U937 cells were transduced with the generated lentivirus and incubated at 37 °C for 48 h. Following incubation, the cells were selected using 2ug/ml puromycin for 2 weeks. Stable FcγRI expression clones were screened using single cell sorting and validated using flow cytometry.

### Viral entry and replication assay

ZIKV entry and replication in U937 cells were performed as previously described^18,69^. Briefly, mAbs were incubated with ZIKV with an MOI of 1 for 1 h at 37 °C to generate mAb:ZIKV immune complexes. The immune complexes and U937 cells were then chilled on ice for 1 h prior to infection. U937 cells were infected with the immune complexes for 1 h on ice and washed 4x with ice cold 1x PBS. Cells were re-suspended in pre-warmed complete RPMI medium and incubated at 37 °C for 2 h to allow virus entry. To assess ZIKV entry, cells were incubated with 0.25% trypsin for 60 min on ice to remove any extracellular virus and washed 4x with ice cold 1x PBS. Cells were then lysed in RNA lysis buffer. For ZIKV replication, cells were incubated for another 12h and 24h after virus entry. Intracellular virus was quantified using qRT-PCR as described above.

### Statistical analysis

Hofbauer cells ADE RT-PCR data was examined using 1-way ANOVA and Tukey’s multiple comparison test, p<0.05. ZIKV entry and replication assay was analyzed by Student’s t-test, p<0.05. Correlations were analyzed using Spearman’s Rho on log-transformed data. All statistical analysis was performed using GraphPad Prism software version 10. p < 0.05 was considered as significant. Results are shown as mean ± SD. Further experimental statistical details can be found in the Figure legends.

## Notes

### Competing Interest Statement

The authors have declared no competing interest.

