## Supplemental File for "FcγRI is the key determinant of antibody-mediated Zika virus infection of human placental macrophages"

### SUPPLEMENTARY MATERIALS

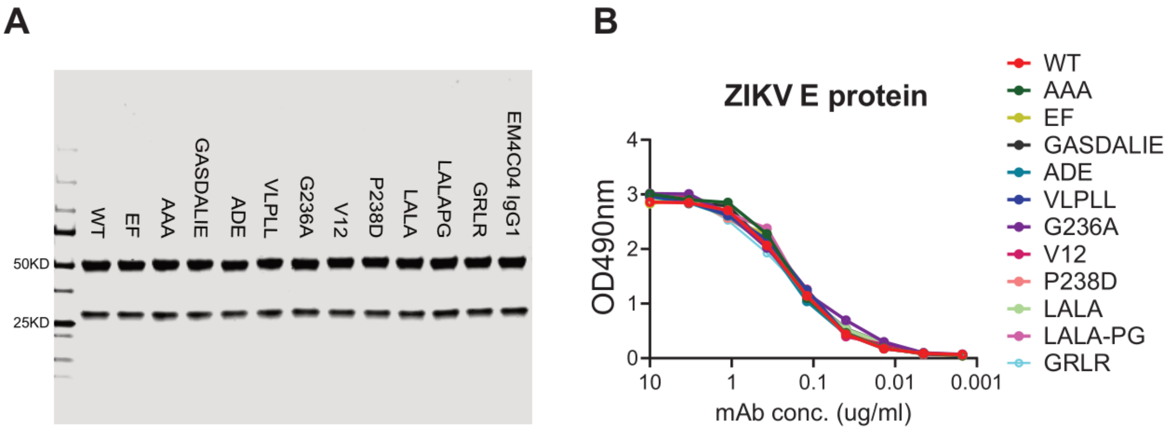


**Figure S1. Expression and characterization of 33.3A06 IgG1 Fc variants.**

**(A)** SDS-PAGE analysis showing purified 33.3A06 WT and Fc variants. EM4C04 IgG1 was an IgG1 control mAb. **(B)** Binding of purified 33.3A06 WT and Fc variants to recombinant ZIKV envelope protein by ELISA. The results plotted are representative of two independent ELISA experiments.

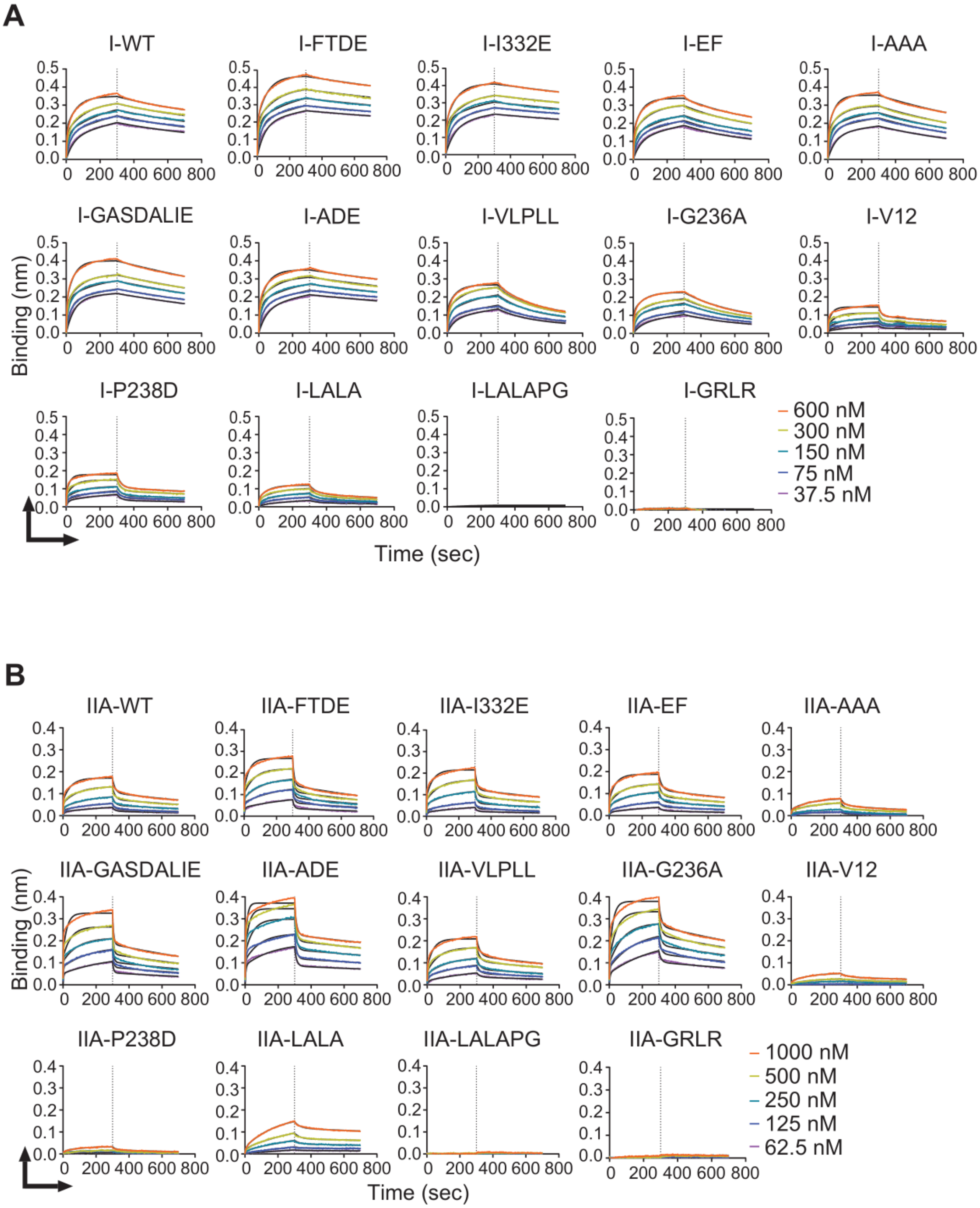
A
I-WT
I-FTDE
I-I332E
I-EF
I-AAA
I-GASDALIE
I-ADE
I-VLPLL
I-G236A
I-V12
I-P238D
I-LALA
I-LALAPG
I-GRLR
Binding (nm)
Time (sec)
600 nM
300 nM
150 nM
75 nM
37.5 nM
B
IIA-WT
IIA-FTDE
IIA-I332E
IIA-EF
IIA-AAA
IIA-GASDALIE
IIA-ADE
IIA-VLPLL
IIA-G236A
IIA-V12
IIA-P238D
IIA-LALA
IIA-LALAPG
IIA-GRLR
Binding (nm)
Time (sec)
1000 nM
500 nM
250 nM
125 nM
62.5 nM

C

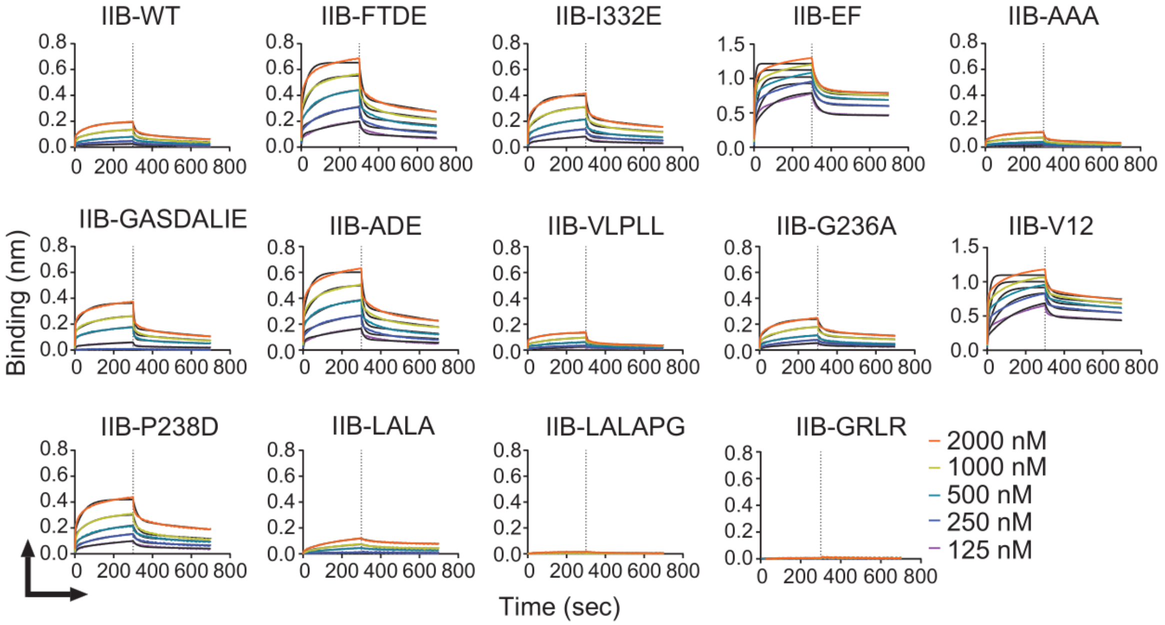
IIB-WT
IIB-FTDE
IIB-I332E
IIB-EF
IIB-AAA
IIB-GASDALIE
IIB-ADE
IIB-VLPLL
IIB-G236A
IIB-V12
IIB-P238D
IIB-LALA
IIB-LALAPG
IIB-GRLR
Binding (nm)
Time (sec)
0 200 400 600 800
2000 nM
1000 nM
500 nM
250 nM
125 nM

D

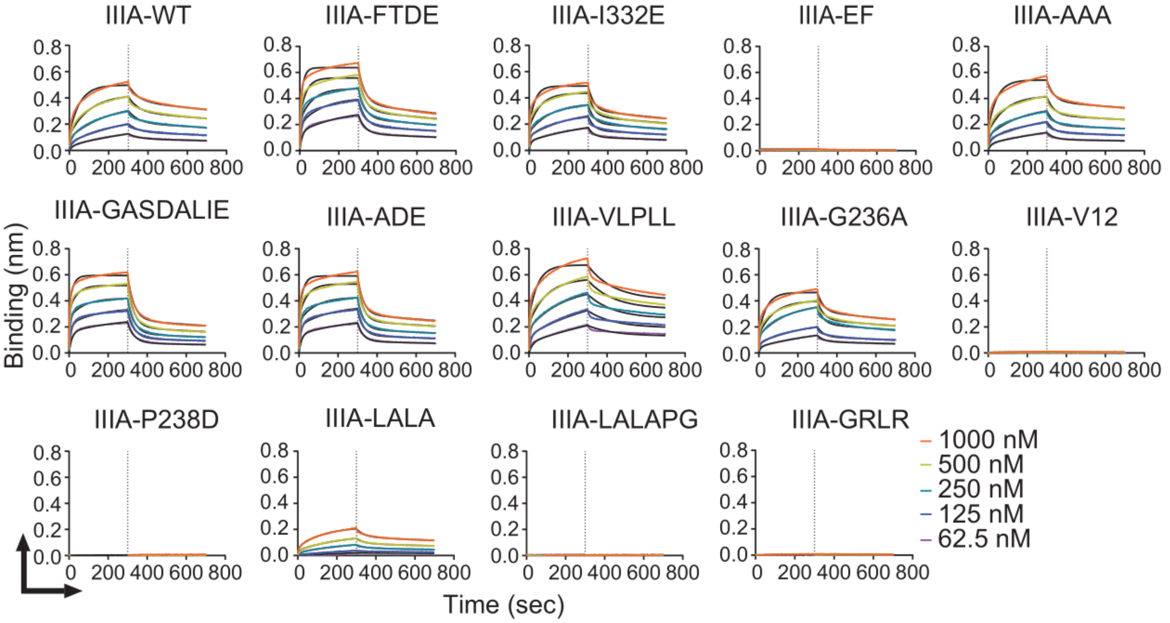
IIIA-WT
IIIA-FTDE
IIIA-I332E
IIIA-EF
IIIA-AAA
IIIA-GASDALIE
IIIA-ADE
IIIA-VLPLL
IIIA-G236A
IIIA-V12
IIIA-P238D
IIIA-LALA
IIIA-LALAPG
IIIA-GRLR
Binding (nm)
Time (sec)
0 200 400 600 800
1000 nM
500 nM
250 nM
125 nM
62.5 nM

**Figure S2. Binding kinetics of 33.3A06 WT IgG1 and Fc variants to human recombinant FcγRs.**

**(A)** BLI curves and fitting curves (in black) obtained for immobilized FcγRI to 33.3A06 WT IgG1 and Fc variants. Antibody concentrations range from 600nM to 37.5nM by 2-fold serial dilution. **(B)** BLI curves and fitting curves (in black) obtained for immobilized FcγRIIA to 33.3A06 WT IgG1 and Fc variants. Antibody concentrations range from 1000nM to 62.5nM by 2-fold serial dilution. **(C)** BLI curves and fitting curves (in black) obtained for immobilized FcγRIIB to 33.3A06 WT IgG1 and Fc variants. Antibody concentrations range from 2000nM to 125nM by 2-fold serial dilution. **(D)** BLI curves and fitting curves (in black) obtained for immobilized FcγRIIIA to 33.3A06 WT IgG1 and Fc variants. Antibody concentrations range from 1000nM to 62.5nM by 2-fold serial dilution. Derived KD values are presented in **Table S1**.

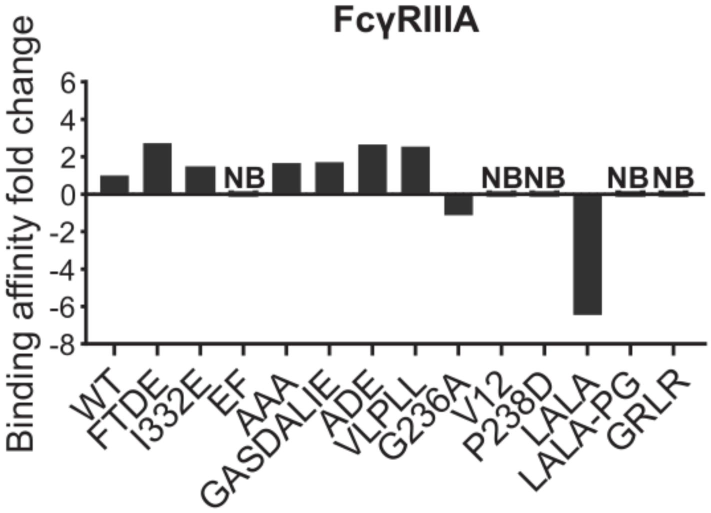


**Figure S3. Fold changes in FcγRIIIA binding affinity of the 33.3A06 Fc variants.** $K_D$ values of the WT IgG1 and Fc variants binding to recombinant human FcγRIIIA were determined by BLI. Relative binding affinity=$K_D$ (WT)/$K_D$ (variant). Variants with negative values show decreased binding affinity. NB, no detectable binding.

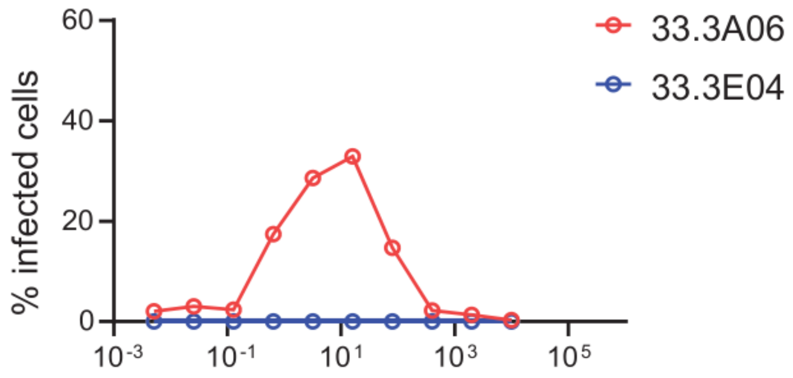


**Figure S4. U937 cells are resistant to ZIKV infection in the absence of cross-reactive antibody.**

U937 cells were infected with ZIKV (MOI 0.5) and in the presence of 33.3A06 or a non-cross-reactive control antibody at a range of mAb concentrations. Infected cells were detected by 4G2 staining at 24 hr post infection (hpi) and flow cytometry. Antibody dilutions were performed in singlicate.

**A**

| mAb | Epitope specificity | Zika virus neutralization IC50 (ug/ml) |
|---|---|---|
| 33.3A06 IgG1 | E dimer | 0.3 |
| 31.3F01 IgG1 | E monomer | >20 |
| 32.2D03 IgG1 | E dimer | 2.5 |
| 32.2E04 IgG1 | E dimer | >20 |
| 33.3F05 IgG1 | E monomer | >20 |
| 39.3D02 IgG1 | E dimer | 20 |
| EDE1 C10 IgG1 | E dimer | 0.1 |
| Z004 IgG1 | EDIII | 0.002 |

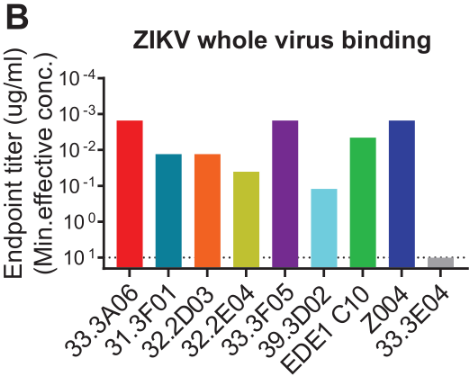


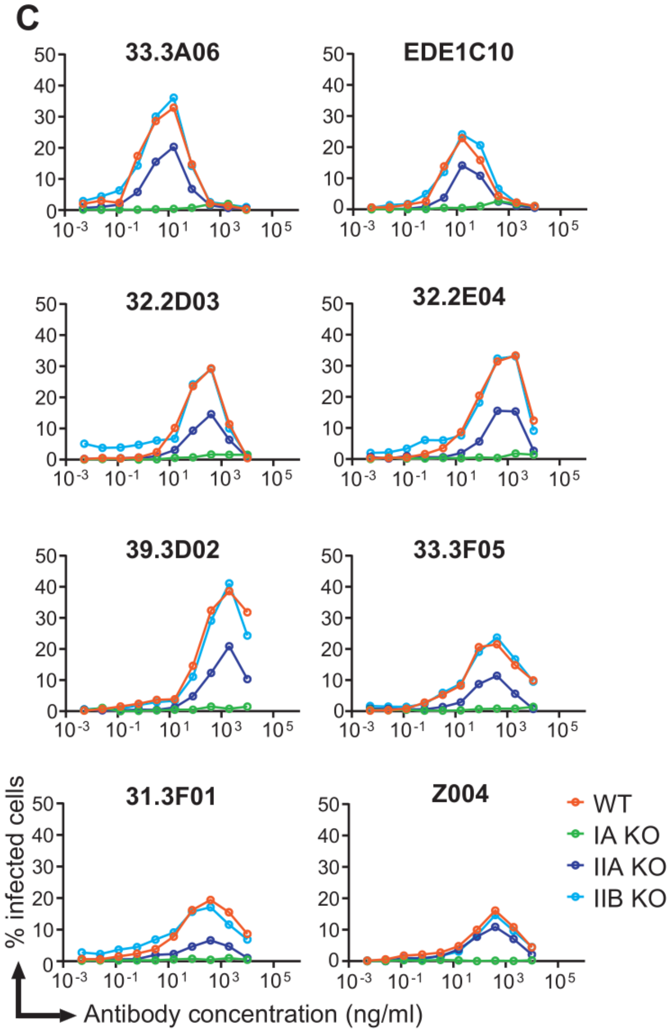


**Figure S5. FcγRI is the primary Fc receptor for antibody-mediated ZIKV infection in U937 cells.**

**(A)** Epitope specificity and neutralization properties of mAbs. **(B)** Binding of mAbs to ZIKV whole virus. Values plotted represent the minimum concentration required three times the background signal of plain blocking buffer. Dotted line represents the maximum mAb concentration used in ELISA: 10 µg/mL. Data shown are representative of three independently performed experiments. **(C)** WT, *FCGR1A* knockout, *FCGR2A* knockout and *FCGR2B* knockout U937 cell lines were infected with ZIKV (MOI 0.5) in the presence of a panel of mAbs. Infected cells were detected by 4G2 staining and flow cytometry at 24 hpi. Data shown are representative of three independently performed experiments. WT, wild type; IA KO, *FCGR1A* knockout; IIA KO, *FCGR2A* knockout; IIB KO, *FCGR2B* knockout.

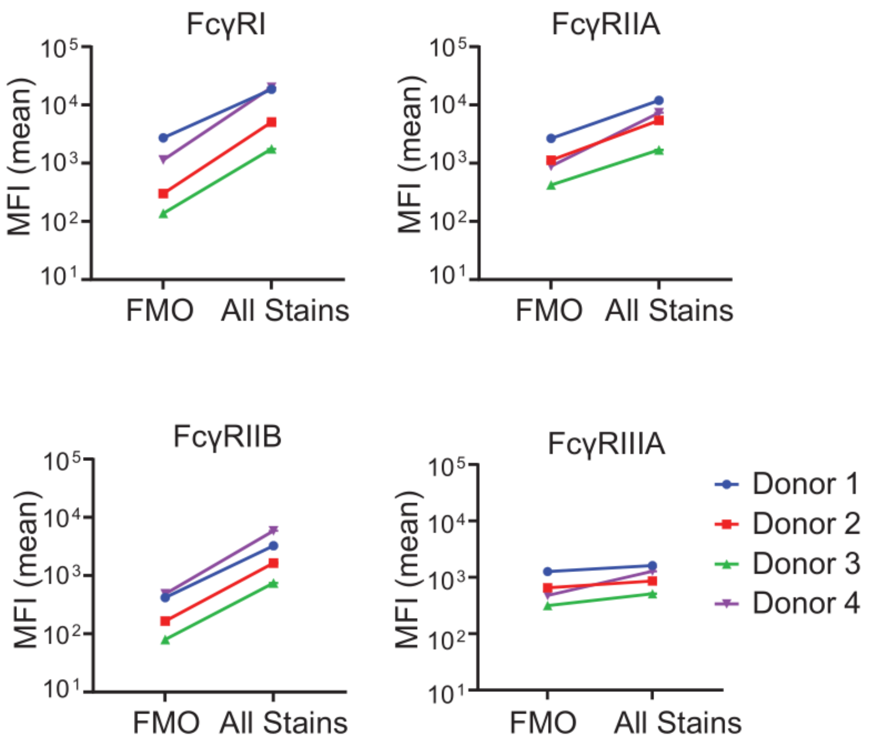


**Figure S6. FcγRs expression on Hofbauer cells.**

Quantification of mean fluorescent intensity (MFI) of FcγRI, FcγRIIA, FcγRIIB and FcγRIIIA expression on Hofbauer cells by flow cytometry. Representative experiments from n = 4 donors.

**Table S1. The list of 33.3A06 IgG1 Fc mutants.** $K_D$ values of the WT IgG1 and Fc variants binding to recombinant human FcγRI, FcγRIIA, FcγRIIB and FcγRIIIA were determined by BLI.

| KD value (nM) | | | | | |
|---|---|---|---|---|---|
| **Fc mutation** | **FcγRI** | **FcγRIIA** | **FcγRIIB** | **FcγRIIIA** | **Ref.** |
| **WT** | **2.7** | **47** | **226** | **25.3** | **-** |
| **H268F/S324T/ S239D/I332E (FTDE)** | **0.52** | **22.9** | **19.8** | **9.26** | (Moore, Chen et al. 2010) |
| **I332E** | **0.86** | **34.3** | **93.8** | **17.5** | (Richards, Karki et al. 2008) |
| **S267E/L328F (EF)** | **3.4** | **36.3** | **0.44** | **NB** | (Chu, Vostiar et al. 2008) |
| **S298A/E333A/K334A (AAA)** | **2.77** | **495** | **673** | **15.2** | (Shields, Namenuk et al. 2001) |
| **G236A/S239D/A330L/I332E (GASDALIE)** | **1.87** | **22** | **141** | **14.8** | (Ahmed, Keremane et al. 2016) |
| **G236A/S239D/I332E (ADE)** | **1.75** | **8.2** | **20.7** | **9.5** | (Richards, Karki et al. 2008) |
| **L235V/F243L/R292P/Y300L/P396L (VLPLL)** | **4.6** | **28.9** | **471** | **9.98** | (Nordstrom, Gorlatov et al. 2011) |
| **G236A** | **8.2** | **15.7** | **162** | **28** | (Richards, Karki et al. 2008) |
| **E233D/G237D/P238D/H268D /P271G/A330R (V12)** | **13.7** | **872** | **1.79** | **NB** | (Mimoto, Katada et al. 2013) |
| **P238D** | **7.9** | **NB** | **69.5** | **NB** | (Mimoto, Katada et al. 2013) |
| **L234A/L235A (LALA)** | **60.9** | **248** | **1070** | **163** | (Xu, Alegre et al. 2000) |
| **L234A/L235A/P329G (LALA-PG)** | **NB** | **NB** | **NB** | **NB** | (Schlothauer, Herter et al. 2016) |
| **G236R/L328R (GRLR)** | **NB** | **NB** | **NB** | **NB** | (Chu, Vostiar et al. 2008) |
